# Behavioral Repertoire of lab-reared early juveniles of the Mexican four-eyed octopus: *Octopus maya*

**DOI:** 10.1101/2024.04.14.589436

**Authors:** D.A. González-Navarrete, F. Vergara-Ovalle, P. García-Andaluz, F. Ayala-Guerrero, C. Rosas, P. Vázquez-León, D.B. Paz-Trejo, H. Sánchez-Castillo

## Abstract

Behavioral studies have predominantly focused on organisms within the phyla Craniata and Arthropoda. Yet, there has been a growing interest in studying the behavior of organisms from alternative phyla, such as mollusks, owing to the research opportunities they offer. Among mollusks, cephalopods have emerged as a prominent subject of inquiry. However, behavioral research on Mexico’s endemic species, *Octopus maya (Om)*, remains conspicuously scarce. *Om* exhibits favorable attributes for utilization as a standardized animal model in neuroscience research, primarily due to its adaptability to laboratory settings and the successful raising of multiple generations. A comprehensive understanding of *Om*’s behavior within laboratory environments is essential to harness its potential as a research model. Thus, the main goal of this study was to establish a comprehensive behavioral catalog for *Om* under laboratory conditions. Thirteen *Om* subjects (6 to 20 grams) were housed in controlled tank environments. Our findings reveal that *Om* exhibits a diverse behavioral repertoire, comprising a minimum of twenty-one distinct behaviors categorized into six behavioral classes. Additionally, *Om* displays discernable diurnal and nocturnal activity patterns, with increased activity levels, altered behavior distributions, and varying activity frequencies predominantly during daylight hours. This expanded knowledge of *Om*’s behavior enhances its suitability as a research model organism.

## INTRODUCTION

Behavior constitutes a multidisciplinary subject of inquiry, attracting interest from various fields, including Neuroscience, Ethology, Ecology, and Psychology. For the behavioral analysis, the ethogram constitutes a fundamental tool, essential for a comprehensive study. Historically, behavior studies have focused mainly on animals of two phyla in particular: Craniata and Arthropoda. However, it is noteworthy that the scope of behavioral research has expanded to encompass a broader range of phyla in recent years (Schnell & Clayton, 2021). Behavioral research within diverse phyla, such as Mollusca, has gained increasing prominence. Among mollusks, cephalopods, including nautilus, squids, cuttlefishes, and octopuses, have emerged as particularly promising subjects for research inquiries (Mather, 2008). The neural arrangement of octopuses affords them a high degree of cognitive complexity, encompassing abilities such as attention, learning, memory, among others, surpassing those observed in other invertebrate organisms (O’Brien et al., 2018; Grasso & Basil, 2009). Their complex nervous system, alongside their efficient physiology and motor control, establishes the basis for complex behavior and allows them to perform a wide behavioral repertoire (Hochner, 2008; Mather, 2008), making a good candidate for a wide range of neuroscience studies (behavioral, ethological, molecular, cellular, etc.).

Comprehensive descriptions of the behaviors exhibited by select octopus species have been documented (Mather & Scheel, 2014; Borrelli et al., 2006; Mather et al., 2010; Hanlon & Messenger, 1996; Wells, 1978). However, it is noteworthy that few studies have provided a comprehensive account of the entire behavioral repertoire exhibited by octopuses. One of those is a specific ethogram of *Abdopus aculeatus* (Huffard, 2007), and another one is a comprehensive overview of general behaviors across seventeen members of the Octopodidae family (Mather & Alupay, 2016). Other articles have described individual behaviors of different octopuses (*Octopus vulgaris*, *Eledone cirrhosa,* and *O. bimaculoides),* among which are the description of components, postures, and actions in the use of arms (Mather, 1998), feeding and foraging behaviors (Mather, 1991), patterns of coloration and movement of the whole body after exposure to an aversive stimulus (Packard & Sanders, 1971), den occupancy, maintenance, and blocking behaviors (Mather, 1994), attack and withdrawal behaviors related to den use (Cigliano, 1993), inactivity, activity and alerting behaviors (Cobb et al., 1995) and sleeping, resting, hunting, feeding, den maintenance, and grooming (Mather, 1988).

The study of neurobiology and behavior in octopuses within their natural habitat poses considerable challenges. Consequently, a predominant approach involves conducting such studies with wild-caught specimens maintained under controlled laboratory conditions. However, this approach involves its own set of problems, primarily stemming from the inherent lack of control over developmental variables, including the life history of the subjects and any prior exposure to aversive stimuli. These variables have the potential to exert unforeseen influences on behavioral performance, resulting in differential responses to identical stimuli. To mitigate the impact of these variables, the use of octopus species that have adapted to laboratory conditions and can be reared in captivity from hatching emerges as a viable and advantageous strategy. These species serve as well-suited candidates for the research of behavior in octopuses, offering an opportunity to establish an optimal reference point for neuroscience research, while simultaneously affording greater control over experimental variables.

In this context, *Om*, (Mexican four-eyed octopus, Figure 1A), emerges as a promising candidate for in-depth investigations into neurobiology and behavior (Vergara-Ovalle et al., 2023; Hanlon & Forsythe, 1985). *Om* has adapted very well to laboratory conditions, has been successfully bred in captivity through several generations (Rosas et al., 2007; Hanlon & Forsythe, 1985; Van Heukelem, 1976). *Om* stands out as a promising species for aquaculture due to its rapid transition from holobenthic hatchlings to benthic juveniles within 7 to 10 days post-hatching (Rosas et al., 2014). Also, significant progress has been made in the *O. maya* aquaculture, exemplified by the development of new outdoor tank designs and the formulation of a successful diet enabling the animals to attain a body weight of 250 g within 120 days after hatching (Sykes et al., 2014; Martinez et al., 2014). Currently, captive breeding in México is being developed by Universidad Nacional Autónoma de México (UNAM) at the Unidad Multidisciplinaria de Docencia e Investigación (UMDI) in Sisal, Yucatán since 2004 (Rosas et al., 2014), providing this species with a great advantage as a research model for octopus’ behavior, reducing the number of variables that could affect the behavioral studies. The main objective of this study was to construct a captive activity budget delineating the behaviors exhibited by juvenile subjects of the species *Om*, which were reared and maintained under controlled laboratory conditions. Plus, having a baseline quantification of *Om* behaviors in the laboratory will allow objectively quantifying changes in behavior, due to variable manipulations in future neurobiological research. The outcome of this work establishes a behavioral standard for forthcoming neuroscience research, facilitating a deeper comprehension of the species’ behaviors and aiding in the advancement of future research. It also provides deeper knowledge about specific octopus behavior, which is fundamental for evolutionary and ecological studies.

**Figure 1.**
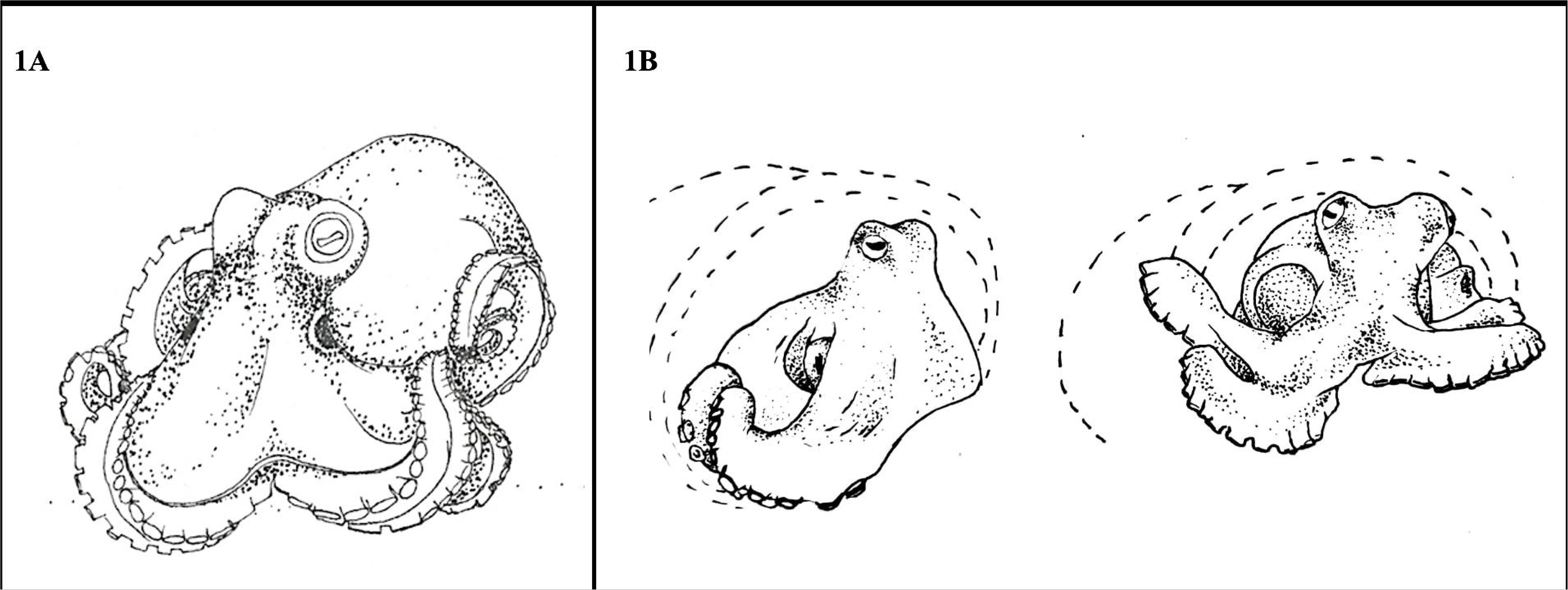
*Octopus maya* illustration (A) and Rest behavior of *Om* (B).

## MATERIALS AND METHODS

### Subjects

13 subjects of *Om* (6-20g) were observed. All octopuses were bred and raised in laboratory conditions. They were obtained from the Applied Ecophysiology Laboratory (UMDI Sisal, UNAM, Mexico). The octopuses were housed in two large tanks of 120cm x 40cm x 40cm (length, width, and height) to keep two specimens per tank separately (each one on one side of the acrylic partition). Weekly water changes and full monthly cleaning procedures of tanks were performed.

Octopuses were maintained in artificial seawater (salinity 3.5%, pH 8, Nitrite 0, Nitrate 25, Ammonia 0, O2 > 95%) in a closed circulation system on a 12-12h light cycle with white and red LED lights. Parameters in tanks were tested every third day during the first 2 weeks of acclimatization and weekly during the rest of the housing time. The subjects were fed with frozen pieces of shrimp or squid tentacles twice a day (10:00 am and 6:00 pm). The health of the subjects was monitored according to the Guidelines for the Care and Welfare of Cephalopods in Research (Fiorito, et al., 2015).

### Behavioral record

Two types of behavioral recordings were performed. First, nine subjects (at different times, over one year) were used for an *ad libitum* sampling of the behavioral repertoire through short videos (5 - 10 minutes). The recordings began any time the octopuses showed a change in activity and continued until the animal completely stopped the behavior. These recordings were used to obtain a catalog of all behaviors presented by the subjects.

Also, four different subjects were simultaneously used to determine periods of activity, frequency, and duration of behaviors displayed, through continuous 24-hour recording without the presence of the researchers (except during feeding periods) and maintaining a 12/12 light cycle. For this, a full HD video camera (Sony, Inc.) was mounted on a tripod 70 cm away from the tanks and connected to a computer for storage and posterior analysis of videos.

### Behavioral analysis

Observed behaviors from the *ad libitum* sampling were identified, categorized, and operationally defined. Behaviors were included into six different behavioral categories (rest, locomotion, feeding, den maintenance, protection, and others) and representative postures of each were illustrated. Postures were considered as a constant body configuration maintained over time, which allows a better understanding of the behavior that the subjects are displaying at a given time.

The previously described behaviors were used to analyze the 24-hour continuous recordings of four subjects. The duration of behaviors was grouped in 10-minute intervals for the analysis to obtain the periods of activity and non-activity through the 24 hours. Also, the frequency of appearance of each behavior was grouped to observe which behaviors occur most frequently throughout the day.

### Statistical analysis

Inter-observer reliability of analyses was obtained using Cohen’s Kappa coefficient. To determine differences in activity periods, the 24 hours were divided into the hours corresponding to the day (12 hours) and the hours corresponding to the night (12 hours).

Besides, a Nonparametric Wilcoxon test was run. Finally, to determine the most frequent behaviors throughout 24 hours of the day, we run a non-parametric test of Friedman’s ANOVA followed by an uncorrected Dunn’s multiple comparison tests of all activity-related behaviors, to identify differences between each of them.

## RESULTS

A value of 0.612 was obtained after applying a Cohen’s kappa coefficient analysis on the obtained videos, which indicates a good level of agreement.

### Description of the behaviors

The inventory of behaviors for *Om* in laboratory conditions was composed of twenty-three behaviors divided into six behavioral categories (Table 1). Due to incomplete descriptions in the literature on the behavior of *Om*, the descriptions were made only in a structural way. We followed the terminology given by previously reported octopus ethograms (Mather & Alupay 2016; Hanlon et al. 1999 Packard & Sanders, 1971). Although the terms for describing each behavior are the same, we observed differences between previous descriptions and the behaviors of *Om.* That is why the definition and an illustrative image representing the characteristic posture of each behavior, are presented.

**Table 1.** Behavioral categories of *O. maya* and their respective behaviors.

| Rest | Locomotion | Target Directed | Home Maintenance | Protective | Others |
| --- | --- | --- | --- | --- | --- |
| Rest | Posterior Jet<br>Anterior Jet<br>Crawling<br>Climbing<br>Dymantic posture with bipedal locomotion | Attack<br>Reach<br>Prepare Food<br>throwing | Burying<br>Manipulate<br>Carry | Isolation<br>Inking<br>Retroflex<br>Pull/Push<br>Flatten<br>Dymantic posture | Defecation<br>Sucker skin removal<br>Stand tall |

### Rest behaviors

Although octopuses present different sleep states (quiet sleep, active sleep, quiet with open pupil, and alert) (Lima et al. 2021), due to the size of the subjects and low light conditions we were unable to determine sleeping periods on *Om*. Rather we established a resting category for any period of inactivity.

#### Rest

The octopus remains in the same place, without conspicuous movements. The distal part of its front arms points to the posterior side. The mantle is relaxed and parallel to the substrate. It can occur with a head bob, meaning that the head moves up and down, allowing the octopus to better focus on objects in its environment. This resting can occur within or outside their den. Most of the time the subjects rested at the entrance of their den with their head and the first third of their arms outside the den, while the mantle and the rest of the arms remained inside. However, there were also occasions when the subject’s body remained entirely within the den. (Fig. 1B; Supplementary 1A [S1A]).

### Locomotion related behaviors

These behaviors were related to activity and characterized by different forms of movement throughout the environment (Fig. 2).

**Figure 2.**
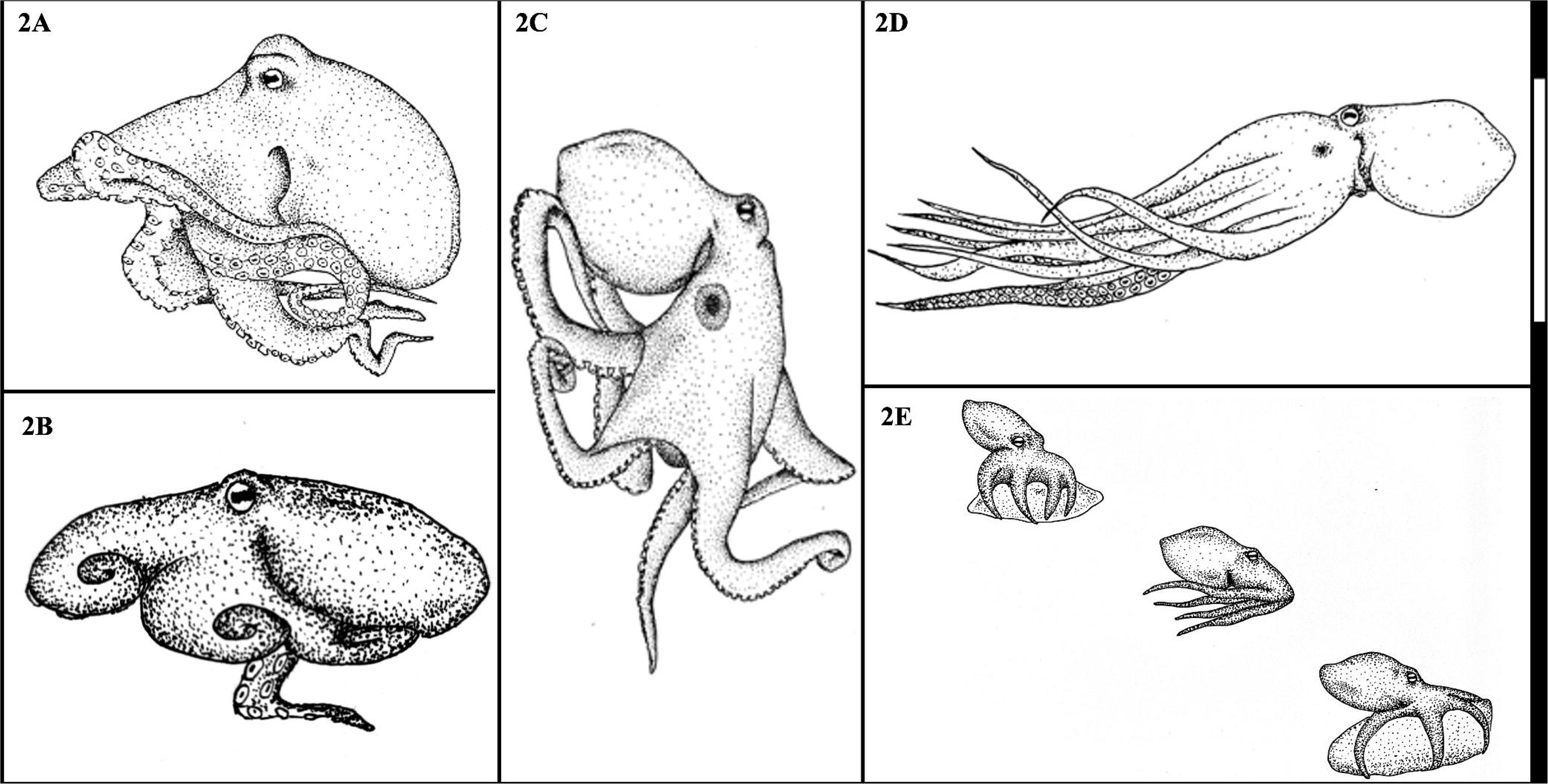
Locomotion behaviors of *O. maya.* A) Crawl: Type of locomotion through the movement of the eight arms together. B) Dymantic posture with bipedal locomotion: The octopus moves with only the fourth and fifth arms. C) Climbing: Locomotion in an ascending and descending way. D) Posterior jet: The octopus makes use of a jet of water to move from one place to another. E) Anterior jet: Locomotion used for jumping on shells or stones, and for hunting.

#### Crawl

This type of locomotion is achieved through the movement of the eight arms together and sequenced. This behavior is the main way of movement for *Om* and occurs when the octopus leaves the den for exploring, disposal wasting, foraging, or mating (Fig A; S1C).

#### Dymantic posture with bipedal locomotion

Also called bipedal walking (Huffard 2007). In this kind of locomotion, the octopus moves through the substrate using only the two more distal arms, while the rest of the arms are partially coiled towards the back of the body and do not touch the bottom. This type of locomotion was observed in short-distance movements and it is possibly a form of crypsis (Hanlon et al. 1999) (Fig 2B; S1D).

#### Climbing

Using its suckers, the octopus moves along the walls of the tank in an ascending, descending, and horizontal way, extending all arms to move. This behavior usually occurs when the octopus is presented with new objects or stimuli; possibly allowing a better visual exploration (see Vergara-Ovalle et al. 2023) (Fig. 2C).

#### Posterior Jet

The octopus makes use of a jet of water through the siphon to move from one place to another faster. In this behavior, the position of the body is similar to other cephalopods, like cuttlefish or squids, since the mantle goes forward, and the arms are in the back. In general, this behavior occurs when the octopus needs to escape or approach a stimulus or a conspecific (Fig. 2D).

#### Anterior jet

The octopus moves in a “jumping” way, using the shells or stones that are close to each other as a platform and positioning itself on top of them. It does that by jetting water in a posterior direction, producing a ventrally directed movement of the animal. The arms and mantle point to the posterior end. Since this type of locomotion can occur while feeding or exploring, sometimes it can be followed by a webover (Fig. 2E; S1F).

### Den maintenance-related behaviors

Behaviors related to the modification and cleaning of the den (Fig. 3).

**Figure 3.**
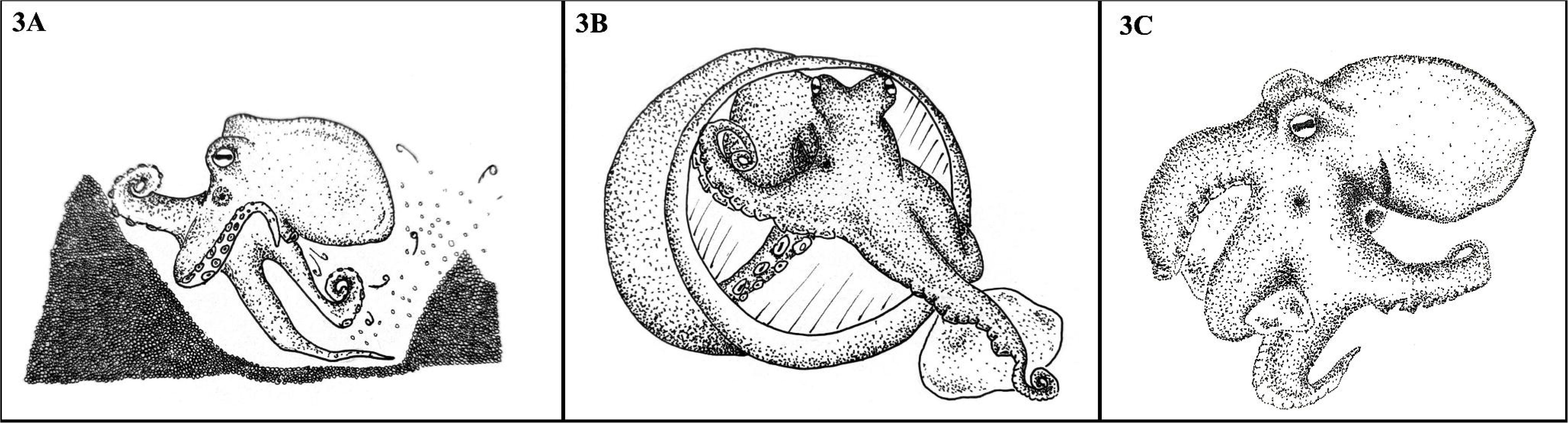
Den maintenance behaviors. A) Burying (Remove + den maintenance throw): The octopus uses the siphon and arms to remove sand under its body. B) Manipulate: The octopus uses one of its arms to hold various objects. C) Carry: The octopus grabs an object and moves with it.

#### Burying (Remove + den maintenance throw)

The octopus uses the siphon to remove sand from under its body. It also uses the back arms to move the sand and stones towards the frontal arms to remove them from the area and make a hole. This behavior is used to create or maintain a den. (Fig. 3A; S1G).

#### Manipulate

The octopus uses one of its arms to hold various objects through the suckers (e.g. shells or stones), either to bring them closer, to move them away, or to rearrange them inside or at the entrance of the den. This can be done by a conveyer belt or by holding with the suckers (see Mather& Alupay 2016) (Fig 3B).

#### Carry

The octopus grabs an object from the environment, such as stones or shells, and holds it with one or two arms and its suckers. Holding said object, it usually moves with crawl-type locomotion. This carry behavior can occur when taking stones or shells to its den or when getting rid of the unconsumed food away from the den (Fig 3C).

### Feeding related behaviors

These behaviors refer to the procedure of getting, ingesting, and eliminating food (Fig 4).

**Figure 4.**
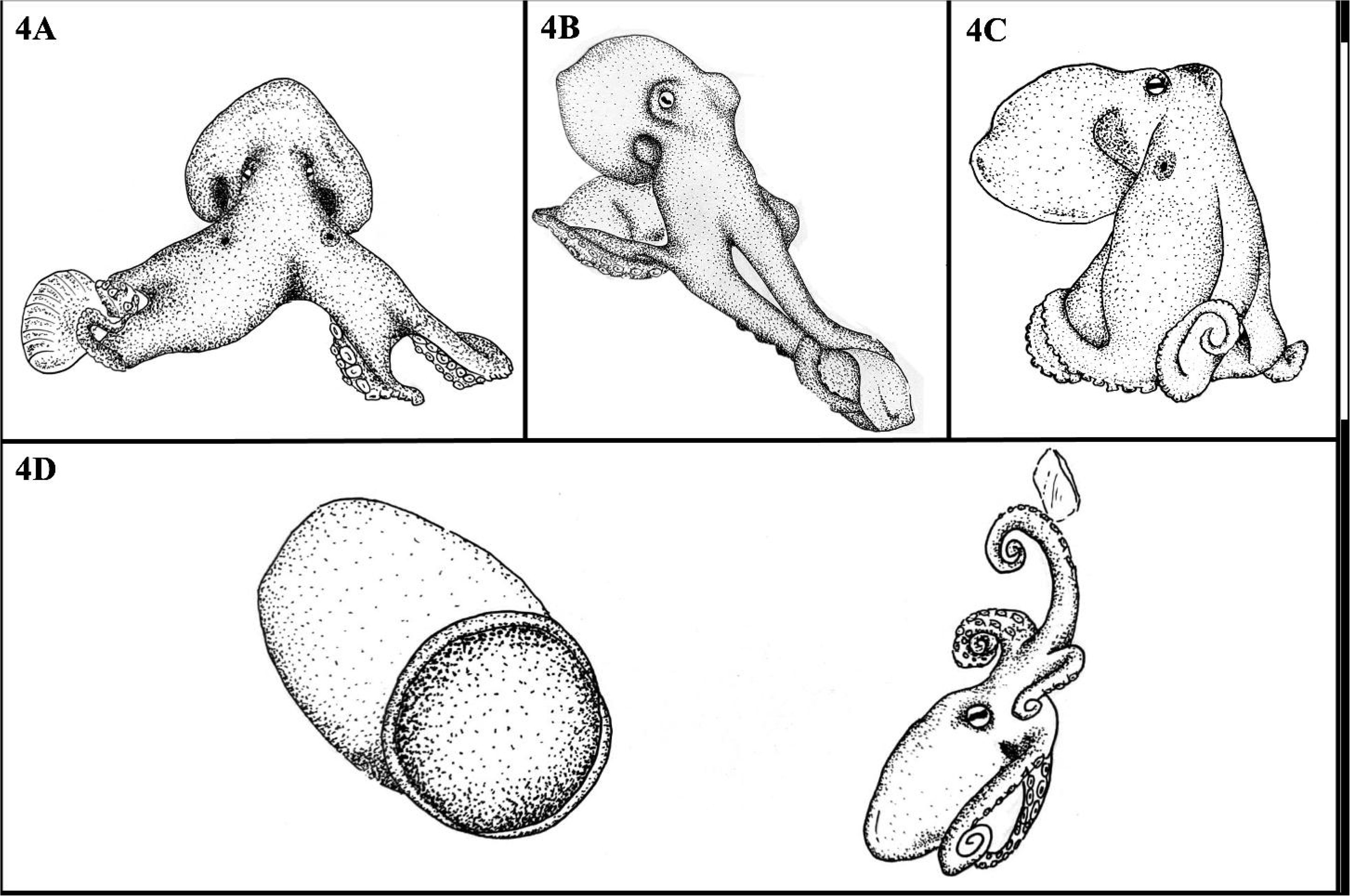
Feeding behaviors. A) Attack: The octopus moves towards the prey with at least two arms and a propulsive movement. B) Reach: The octopus took the food extending and coiling the frontal arms, without moving the rest of the body. C) Prepare: The interbrachial membrane covers the food, causing an elevation of the entire body. D) Food Throwing: The octopus carries uneaten food away from its den.

#### Attack

When the octopus is fed, it moves towards the prey with at least two arms and anterior jetting locomotion, trying to wrap the potential food with the interbrachial membrane (webover), and taking it with the beak. This behavior is most often seen when the octopus is trying to catch living food, however, it can also occur with frozen food (Fig 4A). This behavior can occur during feeding, but has also been observed during foraging, exploration (Young 1956), or social interactions (Tricarico et al. 2011).

#### Reach

The octopus extends one or two of its arms, extending the distal stalks and using the suckers directed towards the object or potential food. After the reach it may happen that the octopus only returns its arm without touching anything, it may only touch the object of interest or it may touch it and grab it with the help of its suction cups. This can occur either in a tip-toe action from the suckers or taking it to the mouth using a conveyer belt movement (see Mather & Alupay 2016) (Fig. 4B; S1E). This behavior can occur during feeding, but also in other contexts such as exploration (Vergara-Ovalle et al., 2023; Gutfreund et al., 1998), social interaction (Scheel et al., 2016) or den maintenance.

#### Prepare

Once the octopus takes the food by displaying some of the previous behaviors (attack; reach), it acquires a specific posture in which the web and the base of the eight arms are covering the food, grabbing the food with the proximal suckers, causing an elevation of the entire body but the distal part of the arms, which are relaxed close to the body. This occurs accompanied by an accelerated contraction and relaxation of the mantle. During this process, the octopus avoids locomotion, staying in one place, most of the time, in its den. (Fig. 4C).

#### Food throwing

When the octopus is done feeding, it carries uneaten food away from its den. For this, it performs a sequence of behaviors that include preparing, carrying the food away from the den, removing (launching a jet of water at high speed through the siphon, by contracting the mantle), and getting back to its den by posterior jetting. (Fig. 4D; S2A). This behavior might be similar to the one described for *O. tetricus* as “debris throwing” (Godfrey-Smith et al. 2022), except that in *O. maya* this behavior seems to be directed towards active waste removal. Also, the behavior may change in the presence of other conspecifics, since our octopuses were isolated in their tanks.

### Protective related behaviors

Behaviors related to the avoidance of aversive stimuli (Fig. 5). These behaviors were only seen during the full cleaning procedure of the tanks.

**Figure 5.**
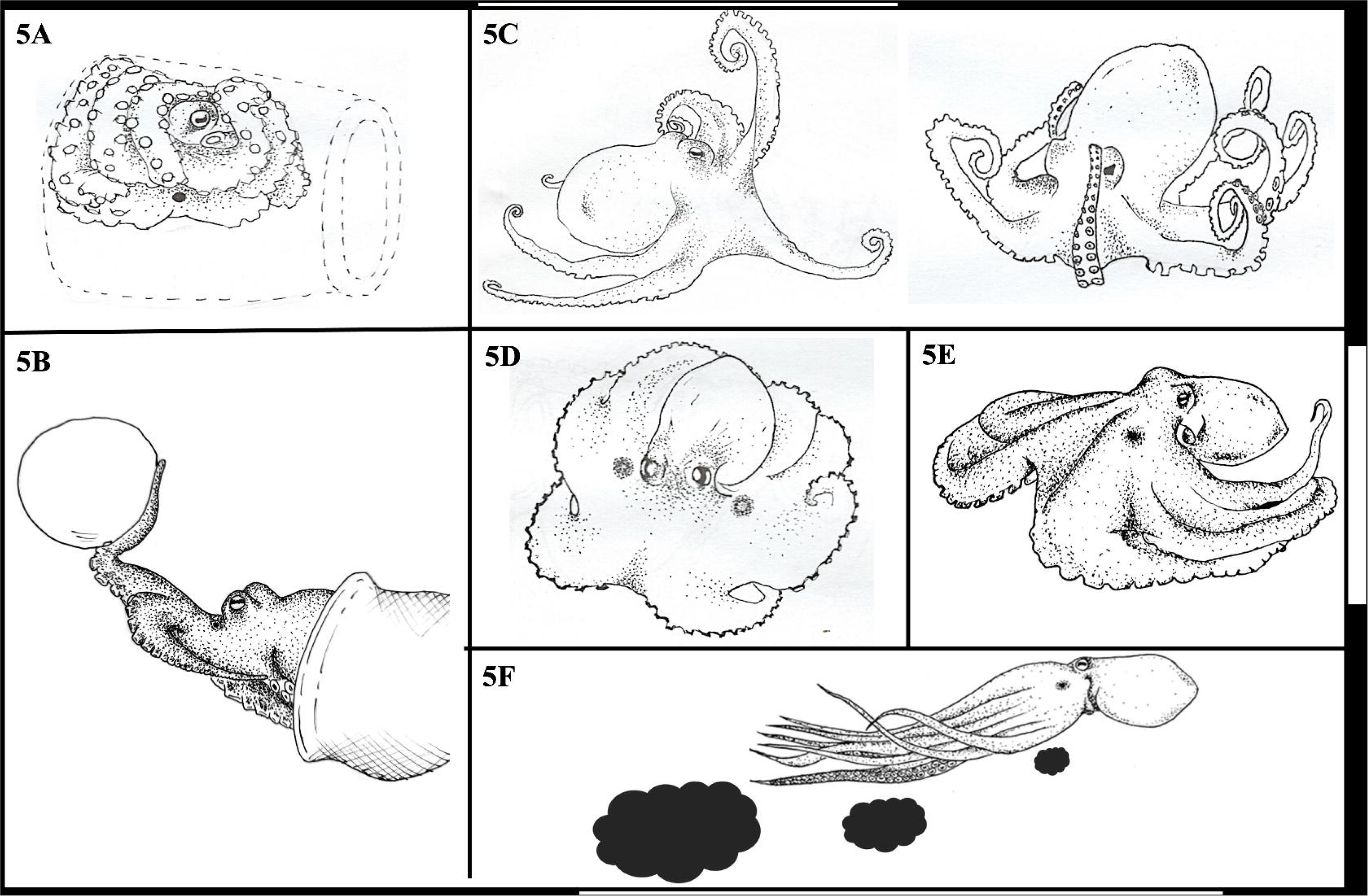
Protective behaviors. A) Isolation: The octopus is completely inside its den and exhibits an anchored posture. B) Pull/Push: The octopus uses one of its arms to move an object closer or away from its body. C) Retroflex: The octopus uses its arms to protect the head and/or mantle from an aversive stimulus. D) Dymantic posture: The octopus’ interbrachial membrane is extended on the surface of the tank. E) Flatten: The entire body of the octopus is closely attached to a surface and spread out, without burrowing into the substrate. F) Inking: The octopus releases a jet of ink through the siphon.

#### Isolation

The octopus’s full body is inside its den and exhibits an anchored posture, characterized by the adherence of the eight arms to the surface of the den, surrounding the mantle and head. Only the beak and one eye are left exposed. This behavior is triggered by an aversive stimulus but is also related to senescence (Fig. 5A).

#### Pull/Push

The octopus uses one of its arms to move an object toward or away from its body. This was observed mainly during feeding, where the octopus repeatedly accepted or rejected food (Fig. 5B; S1H).

#### Retroflex

The octopus arms are bent back toward the posterior part so that the suckers are uniformly presented (Packard & Sanders, 1971). Exposing the front part of the body, partially or totally, by bending the front arms or by curving all of them. This posture might be used to protect the head and/or mantle from an aversive stimulus, but it was also observed in stressed octopuses due to high concentrations of ammonia (data not shown). Similar to the Flamboyant posture in *A. aculeatus* (Huffard 2007) (Fig. 5C; S2B).

#### Dymantic posture

The octopus’ interbrachial membrane or web is completely extended, the base of the arms is slightly arched and the ends are curved in the same direction. The head is slightly elevated and stands out concerning the rest of the body. The chromatophores get contracted producing a pale cream or white color in the animaĺs skin, except for the ocelli, arm crown and the suckers. This behavior is triggered by an aversive stimulus and sometimes is present with locomotion and rest behaviors (Fig. 5D; S2E).

#### Flatten

The entire body of the octopus is closely attached to a surface and spread out, without burrowing into the substrate. The arms can be extended on the surface or retracted close to the body. This behavior is triggered by an aversive stimulus and sometimes is present with locomotion and rest behaviors (Fig. 5E; S2C).

#### Inking

The octopus releases a jet of ink through the siphon, which is triggered by an aversive stimulus (e.g., a predator) and occurs in conjunction with the jet propulsion behavior, allowing a rapid escape movement (Fig. 5F).

### Other behaviors

Behaviors that do not fall into the above categories (Fig. 6).

**Figure 6.**
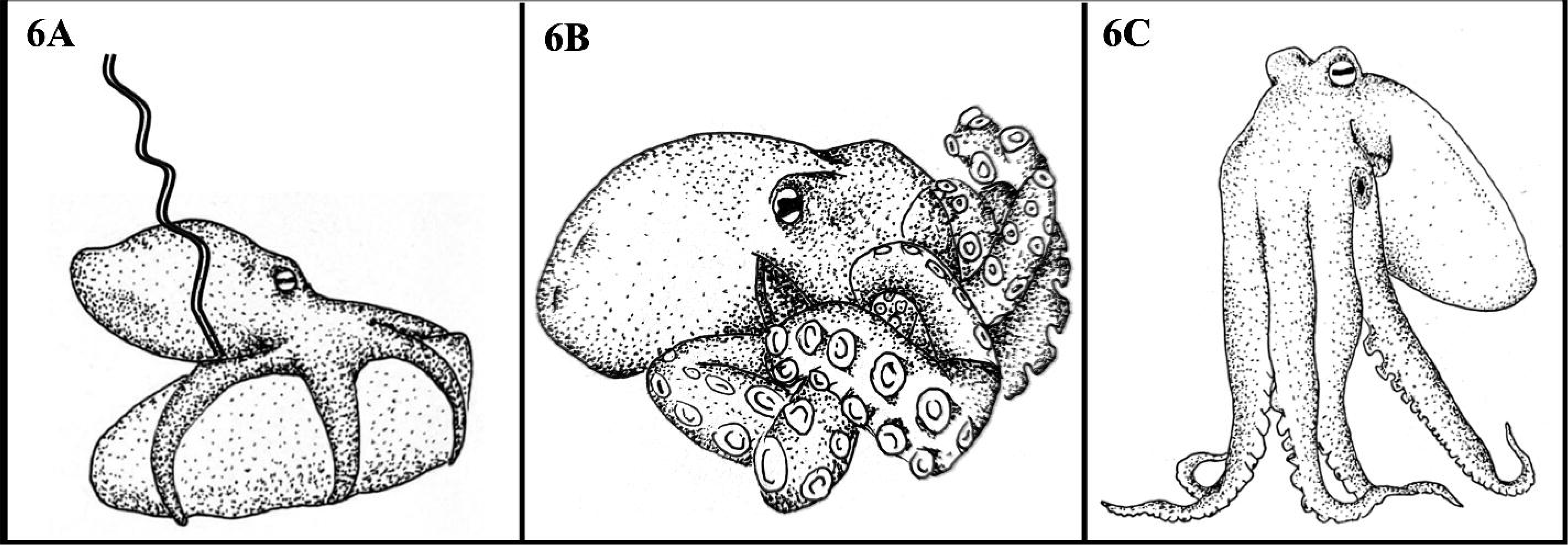
Other behaviors. A) Defecation: Expulsion of organic waste through the siphon. B) Sucker skin removal: The octopus coils all its arms towards the mantle and head, rubbing them in circular movements. C) Stand tall: The octopus puts together the eight arms, extends them, and rests on their tip.

#### Defecation

Expulsion of organic waste through the siphon. This behavior is very quick and does not usually last more than a couple of seconds (Fig. 6A).

#### Sucker skin removal

The octopus coils all its arms towards the mantle and head, rubbing them in circular movements. It’s a behavior related to the cleaning of the octopus itself, such as the elimination of residues, sand, and the layer of chitin that covers the suckers. A type of grooming (Fig. 6B; S2D).

#### Stand tall

The octopus puts together the eight arms, extends them, and rests on their tip, increasing its height and separating the rest of its body from the surface. This behavior can be present at the same time as locomotion and rest behaviors (Fig. 6C).

Moreover, to delineate the activity patterns of *Om*, we conducted a comparative analysis of diurnal and nocturnal activities based on the aggregated 24-hour dataset. We applied the non-parametric Wilcoxon signed-rank test to the combined data from all individuals, yielding statistically significant differences between the day and night activity patterns (p = 0.000602, p < 0.05), as illustrated in Figure 7. These findings indicate distinct periods of activity within the 24-hour cycle. In concert with the distribution of behaviors, our results collectively demonstrate a heightened level of activity during the daytime in contrast to nighttime activity. During the day, the presence of resting behaviors was 80.44%; locomotion behaviors, 10.18%; feeding behaviors, 8.85%; den maintenance behaviors, 0.24%; and other behaviors, 0.30% (Fig. 7). On the other hand, at night the presence of resting behaviors was observed 94.85%; locomotion behaviors, 2.12%; target directed behaviors, 2.58%; den maintenance behaviors, 0.17%; and other behaviors, 0.28%.

**Figure 7.**
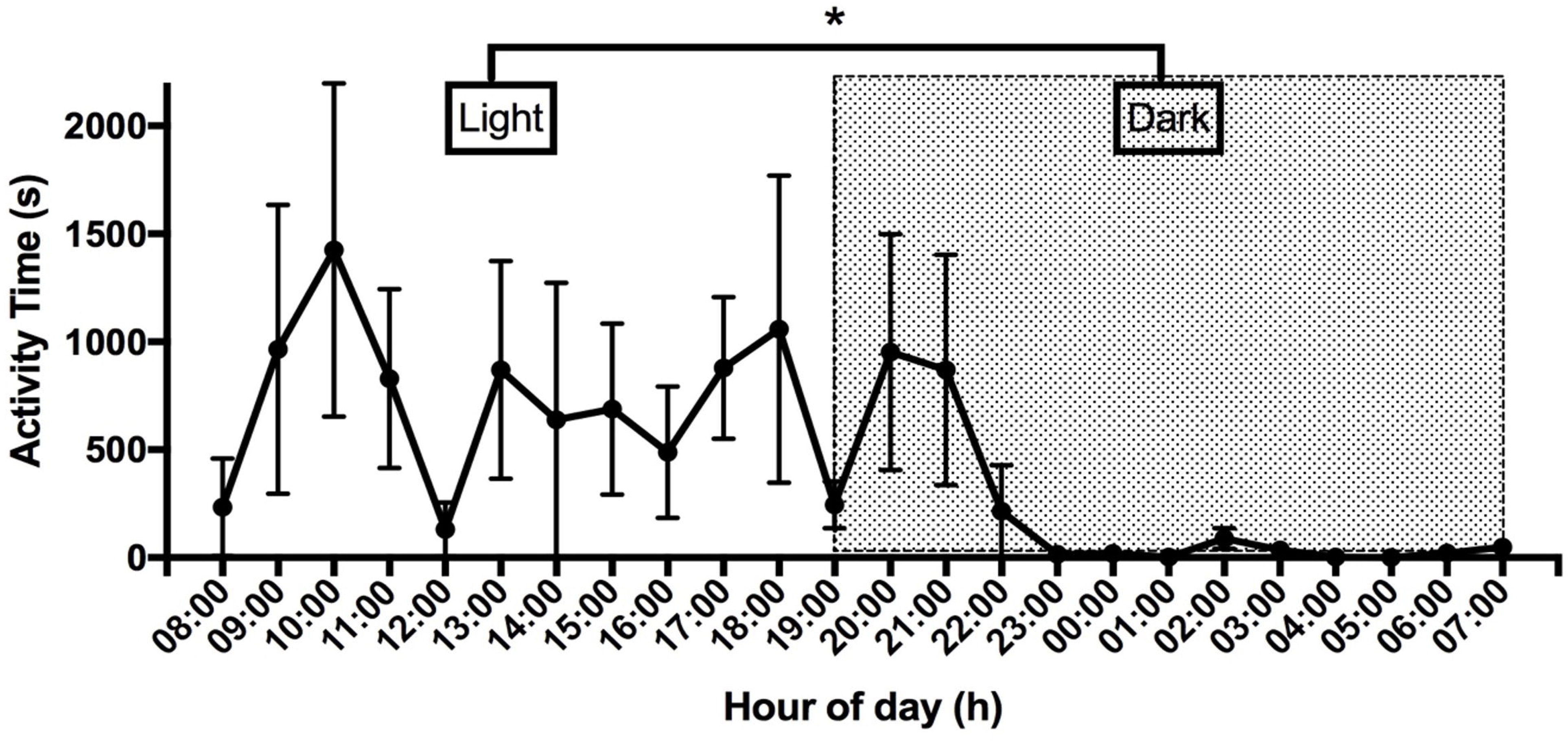
Activity time of the four specimens per hour (24 hours), during the light hours (08:00-19:00), and dark hours (19:01-07:00). Data are presented as mean and standard error. Significance was marked with *(P<0.05).

## DISCUSSION

### Behaviors of *Om*

Literature on octopus behavior often focuses on anecdotal reports of different species, and exhaustive description is quite rare (Marini et al., 2017; Borrelli & Fiorito, 2008). This poses an important problem since the evidence obtained on this model hasn’t had enough experimental control and results vary a lot. In other cephalopod species such as cuttlefish, displayed behaviors have been extensively described, mainly in *Sepia officinalis* (Hanlon & Messenger, 1988) and *Sepia pharaonis* (Nakajima & Ikeda, 2017), being a benchmark for understanding the behavioral ecology of the genus *Sepia* and allowing quantifiable behavioral studies in these organisms (Nakajima & Ikeda, 2017). Just as the ethograms have helped other research models, this detailed behavioral catalog intends to be used as a tool for behavioral quantification in other research endeavors and for them to be replicable in any laboratory. However, it is important to consider that some behaviors would be missing like the ones during other stages of life or in the presence of conspecifics. Furthermore, despite efforts to replicate substrate, den-like structures, and environmental objects reflective of the octopus’ habitat, laboratory tank conditions are inherently limited in providing the full spectrum of enrichment and ecological diversity present in the natural habitat of octopuses. Notwithstanding these constraints, the main objective of this study has been successfully realized, namely, to establish a foundational dataset encompassing the diversity, duration, and frequency of behaviors exhibited by *Om* under controlled laboratory conditions. This dataset serves as a starting point for future research endeavors, particularly in neurobiology and other related disciplines, facilitating comparative investigations and fostering a deeper understanding of octopus behavior.

There are few ethograms of octopus species, that show quantitative data, which makes comparing behavioral repertoires difficult. However, it has been reported that some behaviors displayed by *Om* are also exhibited by other members of the Octopodidae family (Godfrey-Smith et al. 2022; Mather & Alupay, 2016; Huffard, 2007; Hanlon et al. 1999; Packard & Sanders, 1971). The rest category is very similar to the *O. vulgaris* in their wild environment, in which they spend a large percentage of daylight hours inside their den or some other safe place (Mather, 1988). In addition, *O. vulgaris* also presents inactivity periods throughout the day under laboratory conditions, characterized by showing sporadic movements of the arms or body, but always remaining inside the den (Meisel et al., 2011). Similar to what we observed in *Om*, Huffard (2007) also reported resting behaviors in *A. aculeatus* in wild environments, in which the octopus remains at the entrance of the den with its eyes raised and with the front part of the body exposed. However, owing to the constraints of this research, a comprehensive assessment of mantle color and pupil size during periods of den occupancy was not feasible. Therefore, additional research endeavors are warranted to ascertain the occurrence of sleep phases, besides rest, in *Om*. To facilitate the comparison between the behavior of *Om* and other species, the same table prepared for *A. aculeatus* by Huffard (2007) is shown, with the information corresponding to *Om* (Table 2).

**Table 2.**
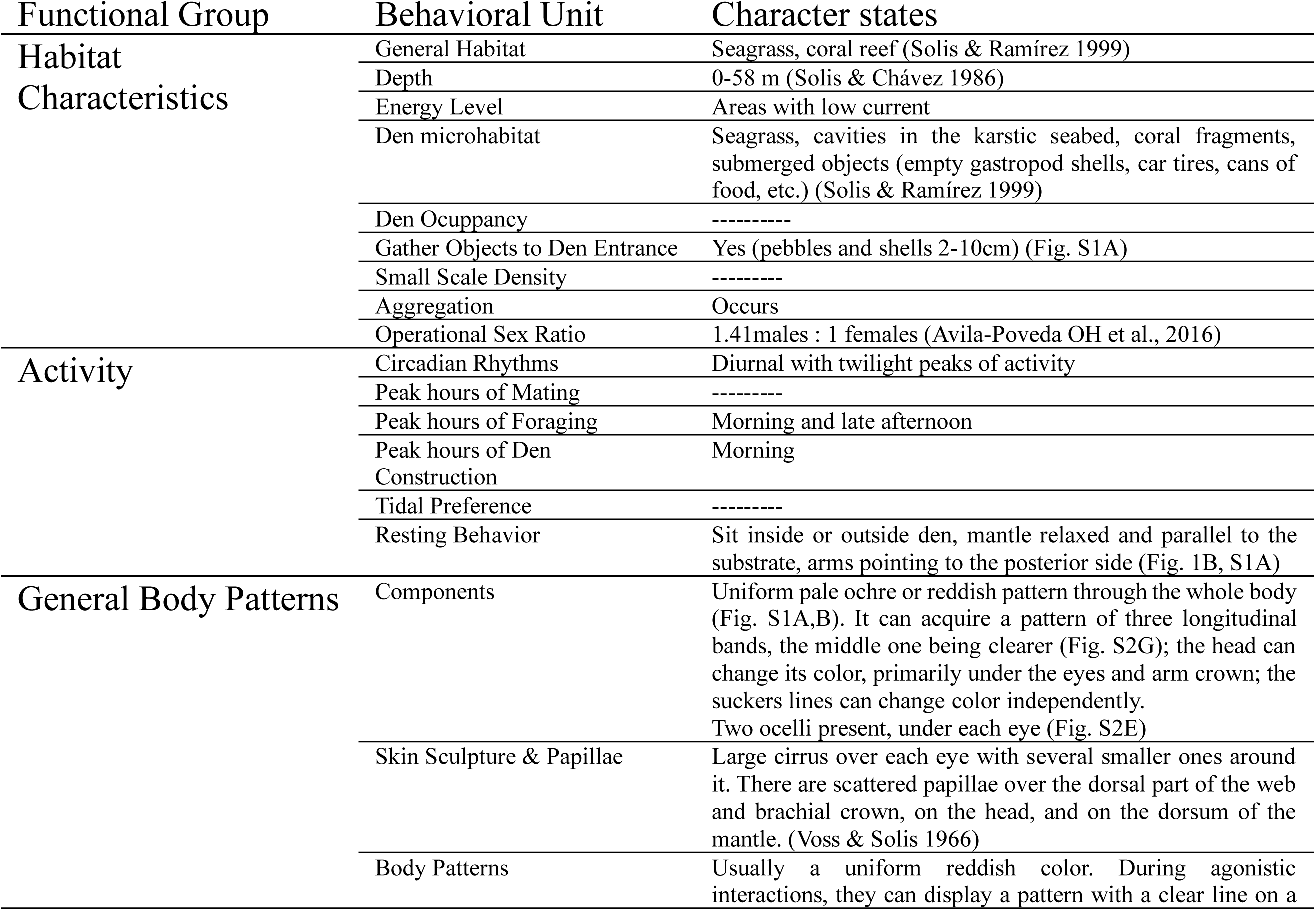
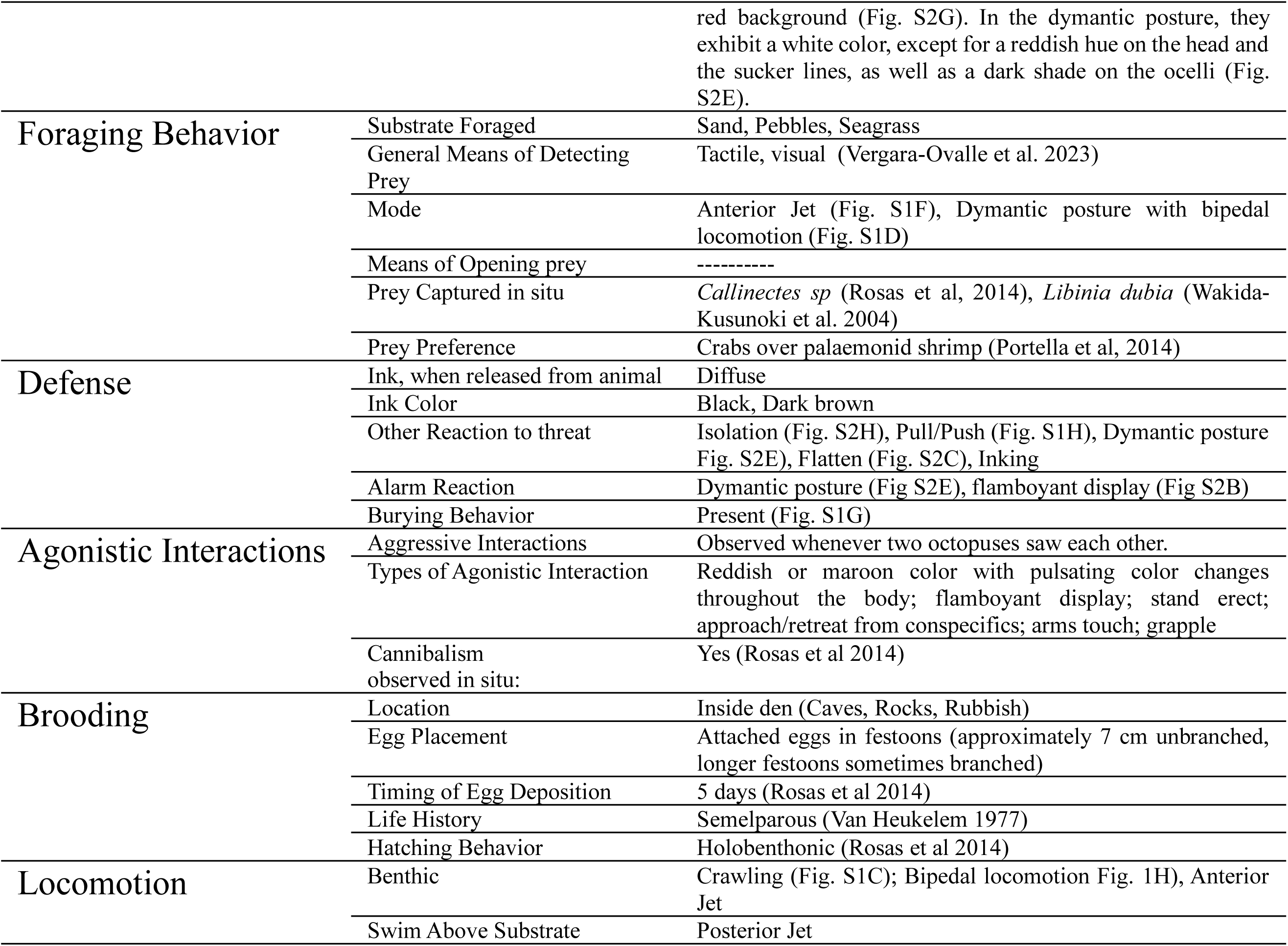
Checklist of behavioral characters for *Octopus maya*.

Regarding locomotion behaviors, *Om* exhibited five distinct forms of locomotion. Crawling is a ubiquitous behavior among benthic octopuses (Mather & Alupay, 2016; Mather, 1998). Nevertheless, variations in crawling behavior may manifest by the specific species. Additionally, the posterior jet behavior, akin to that observed in species such as *A. aculeatus* (Huffard, 2006), *O. vulgaris*, and *O. insularis* (Mather & Alupay, 2016), is characterized by the coordinated movement of the arms held closely together, facilitating rapid relocation. However, it is noteworthy that this behavior can also show subtle differences, with some instances revealing a more pronounced separation between the arms, as observed in *E. dofleini* (Mather & Alupay, 2016). Therefore, the emphasis on a comprehensive investigation of octopus behavior is substantiated by the fact that all known octopus species inhabit diverse marine environments. These varied ecological conditions contribute to the complexity of their central nervous systems and visual pathways (Chung et al., 2022), potentially influencing the observed distinctions in behavioral performance among octopus species.

The process of feeding behavior reported in *Om* is very similar to that described in *O. vulgaris*, both species of octopus have a reaching behavior with their arms towards the food, and both present the same ingestion procedure by transporting the food through the arms towards the interbrachial membrane and maintaining it in the mouth until the feeding is done (Mather, 1991), finally, both also show debris throwing. Likewise, behaviors within the den maintenance category closely resemble those extensively documented in *O. vulgaris* (Mather, 1988; Mather, 1994), involving modifications to the den structure through the removal of substrate material or the placement of rocks and objects at its entrance, similar to observations made in *Om*. In this context, behaviors such as burying and manipulation, aimed at altering object distances and excavating sand, are also consistent with prior reports in *O. vulgaris* (Mather, 1994; Mather, 1998; Mather & Alupay, 2016). Furthermore, behaviors such as sucker skin removal and defecation, as noted in this study, agrees with *O. vulgaris* (Packard & Sanders, 1971; Mather, 1988; Mather & Alupay, 2016), as well as in *A. aculeatus* (Huffard, 2007). Additionally, the behavior of ’standing tall,’ also observed in *Om*, shares similarities with reports of this behavior in *O. vulgaris*, *O. cyanea*, *E. dofleini* (Mather & Alupay, 2016), and *A. aculeatus* (Huffard, 2007).

The retroflex behavior in *Om* was exclusively observed during the cleaning procedure as mentioned above, characterized by the curling of one or all of the arms, a body pattern previously documented in both young specimens (Packard & Sanders, 1971) and adult subjects of *O. vulgaris* (Mather, 1998). In both species, this behavior has been associated with a defensive response to aversive stimuli, such as the presence of potential predators and conspecifics. Moreover, these postures bear resemblance to the ’Flamboyant’ posture, a stereotyped cryptic display characterized by a helical twisting of the arms toward the mantle and head, accompanied by the elevation of the body from the substrate. This posture has been extensively described in young octopus species, including *O. vulgaris*, *E. dofleini, A. aculeatus, O. chierchiae, O. gorgonus, and O. joubini* (Huffard, 2007; Mather & Alupay, 2016; Mather, 1998; Packard & Sanders, 1971).

On the other hand, the inking behavior is one of the most common behaviors in members of the Octopodidae family, so no differences were found in the performance of these behaviors between *Om* and other octopuses (Mather & Alupay, 2016). The flatten behavior has been described in *O. vulgaris, O. cyanea, Enteroctopus dofleini* (Mather & Alupay, 2016), and *A. aculeatus* (Huffard, 2007), and the dymantic posture has also been reported in *O. vulgaris, O. briareus* and *Thaumoctopus mimicus* (Mather & Alupay, 2016; Norman & Hochberg, 2005; Packard & Sanders, 1971).

Finally, regarding the remaining two protective behaviors, namely isolation and pull/push behavior, their occurrence was also confined to the post-cleaning procedure period, specifically when the specimens were reintroduced into their respective individual tanks. Notably, these behaviors exhibited transient characteristics and did not persist over time. Regrettably, the limited availability of studies evaluating the stress responses of octopuses hinders direct comparisons within this behavioral category. Furthermore, it is pertinent to underscore that protective behaviors were exclusively documented during the ad libitum sampling sessions and did not manifest in the analysis of the extended 24-hour video recordings. This observation underscores the adequacy of our laboratory conditions in mitigating behaviors associated with aversive stimuli or environmental stressors. The elucidation of these stress-related behaviors serves as an essential foundation for advancing our understanding of the stress response in octopuses, thereby facilitating future research into the impact of diverse stressors on neurobiology and behavior.

It is noteworthy that the postures and behaviors delineated above have previously been observed in numerous octopus species. However, they may have gone unreported or received limited quantification, primarily due to studies in which behavioral assessments were not the primary focus or because controlled experimental observations were lacking (Marini et al., 2017; Borrelli & Fiorito, 2008). The present behavioral catalog serves a dual purpose: not only does it provide a comprehensive account of behaviors of *Om* in laboratory conditions, but it also addresses the quantification of these behaviors. Therefore, the main objective of this study is to advocate for a heightened focus on behavioral assessments within octopus research. Consequently, it enables the observation of behavioral variations under distinct conditions, facilitating enhanced explorations in the realm of neural system-behavior neuroscience studies.

*Om* demonstrated a substantial allocation of time to resting behavior, constituting 80.44% of their daytime activities and 94.85% during nocturnal periods. This observation aligns with previous empirical evidence suggesting that octopuses, in general, exhibit extended periods of rest throughout the 24-hour day cycle, predominantly within their dens (Mather et al., 1988; Mather, 1985). *Om* exhibited a notable diurnal preference for heightened activity, with a substantial presence of activity-related behaviors predominantly occurring during daylight hours (including locomotion, den maintenance, feeding, and others), distributed between 08:00 and 17:00 hrs. It is important to note that the delineation of activity periods in octopus species remains a complex and evolving area of research, characterized by significant interspecies variation. Furthermore, the precise environmental cues that synchronize octopus activity remain incompletely understood (Mather, 1988). Intriguingly, there have been reports documenting disparate activity periods within the same octopus species (Meisel et al., 2011). For instance, *O. vulgaris* has been observed with both nocturnal activity patterns (Kayes, 1974) and diurnal activity patterns (Mather, 1988). In the case of juvenile *O. vulgaris*, a twilight activity pattern has also been reported (Mather, 1991). The observed variability in octopus activity patterns may be attributed to the influence of environmental factors on the biological rhythms of marine organisms. Variables such as temperature, light levels, and tidal patterns fluctuate across oceanic regions and are likely to contribute to the divergence in reported activity patterns (Ikeda & Yanagisawa, 2018). More research is needed considering these variables to know in depth the origin and maintenance of activity patterns in *Om*.

Based on the findings of this study, *Om* possesses attributes that make it an attractive candidate for research endeavors, owing in part to the successful captive breeding initiatives undertaken in Mexico since 2004, facilitated by the Universidad Nacional Autónoma de México (UNAM). Additionally, the species’ adaptability to laboratory conditions offers a distinct advantage for its utilization in neuroscience research. The ability to maintain *Om* in laboratory settings over extended periods, starting from hatching, is facilitated by the fact that the embryos hatch as holobenthic juveniles, sharing gross anatomical features with adult counterparts (Vidal et al., 2014; Moguel et al., 2010; Rosas et al., 2007; Hanlon & Forsythe, 1985).

The findings of this study reveal that *Om* exhibits behaviors comparable to those observed in other octopus species. Furthermore, a comprehensive understanding of the behavioral repertoire of this species provides a valuable foundation for subsequent investigations of behavior. These results serve as a foundational resource for identifying quantifiable behaviors in future research involving *Om.* Furthermore, they provide insights into the normal daily activity patterns of this species, facilitating the detection of deviations from these patterns and their potential correlation with environmental variables or stressors. Consequently, this study establishes a precedent for future behavioral research endeavors, particularly in the domains of neuroscience and cognition, focusing on *Om*.

## CONFLICTS OF INTERESTS

There are no conflicts of interest to declare.

All applicable international, national, and/or institutional guidelines for the care and use of animals were followed.

## ACKNOWLEDGEMENT

Authoŕs would like to thank M.Sc. Ignacio Morales-Salas from the Aquarium of the Faculty of Sciences, UNAM for his support in the care and maintenance of the specimens.

## REFERENCES

Avila-Poveda OH, Noussithe K, Villalobos B, Santos-Valencia F, Benitez J, Rosas C (2016) Reproductive traits of Octopus maya (Cephalopoda: Octopoda) with implications for fisheries management. Molluscan Research. 36. 29–44. 10.1080/13235818.2015.1072912.

Borrelli L, & Fiorito G (2008) Behavioral Analysis of Learning and Memory in Cephalopods. In Byrne, J.J. (Ed.) Learning and Memory: A Comprehensive Reference. 605–627. 10.1016/B978-012370509-9.00069-3

Borrelli L, Chiandetti C, Fiorito G (2020) A standardized battery of tests to measure Octopus vulgaris’ behavioural performance. Invertebrate Neuroscience, 20(1), 1–15. 10.1007/s1015 8-020-0237-7

Borrelli L, Gherardi F, Fiorito G (2006) A catalogue of body patterning in Cephalopoda. Firenze University Press.

Chung WS, Kurniawan N, Marshall J (2022) Comparative brain structure and visual processing in octopus from different habitats. Current Biology, 32(1), 97–110.e4. 10.1016/j.cub.2021.10.070

Cigliano J (1993) Dominance and den use in *Octopus bimaculoides*. Animal Behaviour, 46 (4), 677–684.

Cobb C, Pope S, Williamson R (1995) Circadian rhythms to light-dark cycles in the lesser octopus, *Eledone cirrhosa*. Marine and Freshwater Behaviour and Physiology, 26:1, 47–57. 10.1080/10236249509378927

Di Cosmo A, Maselli V, Polese G (2018) *Octopus vulgaris*: An Alternative in Evolution. In M. Klock & J. Z. Kubiak (Eds.), Marine Organisms as Model Systems in Biology and Medicine (pp. 585–598). Springer. 10.1007/978-3-319-92486-1_26

Fiorito G, Affuso A, Anderson DB, Basil J, Bonnaud L, Botta G, Cole A, D’Angelo L, De Girolamo P, Dennison N, Dickel L, Di Cosmo A, Di Cristo C, Gestal C, Fonseca R, Grasso F, Kristiansen T, Kuba M, Maffucci F, … Andrews P (2014) Cephalopods in neuroscience: regulations, research and the 3Rs. Invertebrate Neuroscience, 14(1), 13–36. 10.1007/s10158-013-0165-x

Fiorito G, Affuso A, Basil J, Cole A, De Girolamo P, D’Angelo L, Dickel L, Gestal C, Grasso F, Kuba M, Mark F, Melillo D, Osorio D, Perkins K, Ponte G, Shashar N, Smith D, Smith J, Andrews PL (2015) Guidelines for the Care and Welfare of Cephalopods in Research -A consensus based on an initiative by CephRes, FELASA and the Boyd Group. Laboratory Animals, 49 (S2), 1–90. 10.1177/0023677215580006

Godfrey-Smith P, Scheel D, Chancellor S, Linquist S, Lawrence M (2022) In the line of fire: Debris throwing by wild octopuses. PLOS ONE, 17(11), e0276482–e0276482. 10.1371/journal.pone.0276482

Grasso F & Basil JA (2009) The Evolution of Flexible Behavioral Repertoires in Cephalopod Molluscs. Brain Behavior and Evolution, 74(1), 231–245. 10.1159/000258669

Gutfreund Y, Flash T, Fiorito G, Binyamin H (1998) Patterns of Arm Muscle Activation Involved in Octopus Reaching Movements. The Journal of Neuroscience, 18(15), 5976– 5987. 10.1523/jneurosci.18-15-05976.1998

Hanlon R & Messenger JB (1996) Cephalopod behaviour. Cambridge University Press.

Hanlon R & Messenger, JB (1988) Adaptive coloration in young cuttlefish (Sepia officinalis L.): the morphology and development of body patterns and their relation to behaviour. Philosophical Transactions of the Royal Society of London. B, Biological Sciences, 320(1200), 437–487.

Hanlon R, Forsythe JM, Joneschild D (1999) Crypsis, conspicuousness, mimicry and polyphenism as antipredator defences of foraging octopuses on Indo-Pacific coral reefs, with a method of quantifying crypsis from video tapes. Biological Journal of the Linnean Society, 66(1), 1–22. 10.1111/j.1095-8312.1999.tb01914.x

Hanlon R, Forsythe JM (1985) Advances in the laboratory culture of Octopuses for biomedical research. Laboratory Animal Science, 35(1), 33–40.

Hochner B (2008) Octopuses. Current Biology, 18(19), R897–R898. 10.1016/j.cub.2008.07.057

Huffard C (2006) Locomotion by Abdopus aculeatus (Cephalopoda: Octopodidae): walking the line between primary and secondary defenses. The Journal of Experimental Biology, 209(Pt 19), 3697–3707. 10.1242/jeb.02435

Huffard C (2007) Ethogram of Abdopus aculeatus (d’Orbigny, 1834) (Cephalopoda: Octopodidae): Can behavioral characters inform octopodid taxonomy and systematics? The Journal of Molluscan Studies, 73(2), 185–193. 10.1093/mollus/eym015

Ikeda Y & Yanagisawa R (2018) Activity rhythms of the shallow-water octopuses *Octopus laqueus* and *Abdopus aculeatus* with special reference to its relation to light and tidal cycles. Biological Rhythm Research, 4(1), 566–580. 10.1080/09291016.2017.1386897

Kayes RJ (1974) The daily activity pattern of *Octopus vulgaris* in a natural habitat. Marine Behavior and Physiology, 2(1), 337–343. 10.1080/10236247309386935

Lehner P (1996) Handbook of ethological methods (2nd ed.). Cambridge University Press.

Malham SK, Lacoste A, Gélébart F, Cueff A, Poulet S (2002) A first insight into stress-induced neuroendocrine and immune changes in the octopus *Eledone cirrhosa*. Aquatic Living Resources, 15(3), 187–192. 10.1016/S0990-7440(02)01173-7

Marini G, De Sio F, Ponte G, Fiorito G (2017) Behavioral Analysis of Learning and Memory in Cephalopods. Learning and Memory: A Comprehensive Reference. 441–462. 10.1016/B978-0-12-809324-5.21024-9

Martínez R, Gallardo P, Pascual C, Navarro J, Sánchez A, Caamal-Monsreal C, Rosas C (2014) Growth, survival and physiological condition of Octopus maya when fed a successful formulated diet. Aquaculture, 426-427, 310–317. 10.1016/j.aquaculture.2014.02.005

Mather JA (1988) Daytime activity of juvenile *Octopus vulgaris* in Bermuda. Malacologia, 29(1), 69–76. https://archive.org/stream/cbarchive_53203_daytimeactivityofuvenile1962/daytimeactivit yofuvenile1962_djvu.txt

Mather JA (1991) Foraging, feeding and prey remains in middens of juvenile *Octopus vulgaris* (Mollusca: Cephalopoda). Journal of Zoology, 224(1), 27–39. 10.1111/j.1469-7998.1991.tb04786.x

Mather JA (1991) Foraging, feeding and prey remains in middens of juvenile *Octopus vulgaris* (Mollusca: Cephalopoda). Journal of Zoology, 224(1), 27–39.

Mather JA (1994) ‘Home’ choice and modification by juvenile *Octopus vulgaris* (Mollusca: Cephalopoda): specialized intelligence and tool use? The Zoological Society of London, 233(1), 359–368. 10.1111/j.1469-7998.1994.tb05270.x

Mather JA (1998) How do octopuses use their arms? Journal of Comparative Psychology, 112(3), 306–316. 10.1037/0735-7036.112.3.306

Mather JA (2008) Introduction to the symposium “Cephalopods: A behavioral perspective”. American Malacological Bulletin, 24(1), 1. 10.4003/0740-2783-24.1.1

Mather JA & Alupay JS (2016) An Ethogram for Benthic Octopods (*Cephalopoda: Octopodidae*). Journal of Comparative Psychology, 130(2), 109–127. 10.1037/com0000025

Mather JA & Scheel D (2014) Behaviour. In J. Iglesias, L. Fuentes & R. Villanueva (Eds.), Cephalopod Culture (pp. 17–39). Springer. 10.1007/978-94-017-8648-5_2

Mather JA, Anderson RC, Wood JB (2010) Octopus: the ocean’s intelligent invertebrate. Timber Press.

Mather JA, Resler S, Cosgrove J (1985) Activity and movement patterns of *Octopus dofleini*. Mar. Behav. Physiol., 11(1), 301–314.

Lima S, Matias M, Henrique Lopes P, Blanco W, Lima FD, Bruno J, Inácio Gomes Medeiros, Sequerra E, Silva Leite T, Ribeiro S (2021) Cyclic alternation of quiet and active sleep states in the octopus. IScience, 24(4), 102223–102223. 10.1016/j.isci.2021.102223

Meisel DV, Byrne RA, Kuba M, Mather JA (2011) Behavioural sleep in *Octopus vulgaris*. Vie et Milieu, 61(4), 185–190.

Moguel C, Mascaró M, Avila-Poveda O, Caamal C, Sánchez A, Pascual C, Rosas, C (2010) Morphological, physiological, and behavioral changes during post-hatching development of *Octopus maya* (Mollusca:Cephalopoda) with special focus on digestive system. *Aquat*. Biol. 9, 35–48.

Nakajima R & Ikeda J (2017) A catalog of the chromatic, postural and locomotor behaviors of the pharaoh cuttlefish (Sepia pharaonis) from Okinawa Island, Japan. Marine Biodiversity, 47(3), 735–753. 10.1007/s12526-017-0649-8

Norman H & Hochberg F (2005) The “mimic Octopus” (*Thaumoctopus mimicus* n. gen. et sp.), a new octopus from the tropical Indo-West Pacific (Cephalopoda: Octopodidae). Molluscan Research, 25(1), 57–70.

O’Brien CE, Ponte G, Fiorito G (2018) Octopus. Reference Module in Life Sciences, 1–7. 10.1016/B978-0-12-809633-8.90074-8

Packard A & Sanders, G (1971) Body patterns of Octopus vulgaris and maturation of the response to disturbance. Animal Behaviour, 19(4), 780–790. 10.1016/S0003-3472(71)80181-1

Rosas C, Cuzón G, Pascual C, Gaxiola G, Chay D, López N, Maldonado T, Domingues P (2007) Energy balance of Octopus maya fed crab or an artificial diet. Marine Biology, 152(2), 371–381. 10.1007/s00227-007-0692-2

Rosas C, Gallardo P, Mascaró M, Caamal-Monsreal C, Pascual C (2014) Octopus maya. En J. Iglesias, L. Fuentes & R. Villanueva (Eds.), Cephalopod Culture (pp. 383–396). Springer. 10.1007/978-94-017-8648-5_20

Rosas C, Mascaró M, Mena R, Caamal-Monsreal C, Domingues P (2014) Effects of Different Prey and Rearing Densities on Growth and Survival of Octopus Maya Hatchlings. Fisheries and Aquaculture Journal. https://hal.science/hal-03639777

Scheel D, Godfrey-Smith P, Lawrence M (2016) Signal Use by Octopuses in Agonistic Interactions. Current Biology, 26(3), 377–382. 10.1016/j.cub.2015.12.033

Schnell AK & Clayton NS (2021) Cephalopods: Ambassadors for rethinking cognition. Biochemical and Biophysical Research Communications, 564(1), 27–36. 10.1016/j.bbrc.2020.12.062

Sykes A, Koueta N & Rosas C. (2014) Historical Review of Cephalopods Culture. 10.1007/978-94-017-8648-5_4

Tricarico E, Borrelli L, Gherardi F, Fiorito G (2011) I Know My Neighbour: Individual Recognition in Octopus vulgaris. PLOS ONE, 6(4), e18710–e18710. 10.1371/journal.pone.0018710

Van Heukelem WF (1976) Growth, bioenergetics and life span of *Octopus cyanea* and *Octopus maya*. PhD Thesis University of Hawaii.

Van Heukelem WF (1977) Laboratory maintenance, breeding, rearing, and biomedical research potential of the Yucatan octopus (Octopus maya). Laboratory animal science, *27*(5 Pt 2), 852–859.

Vergara-Ovalle F, Ayala-Guerrero F, Rosas C, Sánchez-Castillo H (2023) Novel object recognition in Octopus maya. Animal Cognition, 26(3), 1065–1072. 10.1007/s10071-023-01753-6

Vidal EAG, Villanueva R, Andrade JP, Gleadall IG, Iglesias J, Koueta N, Rosas C, Segawa S, Grasse B, Franco-Santos RM, Albertin CB, Caamal-Monsreal C, Chimal ME, Edsinger-Gonzales E, Gallardo P, Le Pabic C, Pascual C, Roumbedakis K, Wood J (2014) Cephalopod Culture: Current Status of Main Biological Models and Research Priorities. Advances in Marine Biology, 67, 1–98. 10.1016/B978-0-12-800287-2.00001-9

Wakida-Kusunoki A, Sansores R, Ramírez M, Rosas R, Cervera K, Mendez J, Aguilar R (2004) Analysis of red octopus *Octopus maya* abundance in Peninsula of Yucatan. Core.ac.uk. oai:generic.eprints.org:13657/core331

Wells MJ (1978) Octopus: the physiology and behaviour of an advanced invertebrate. Chapman & Hall, London.

Young JZ (1956) Visual Responses by Octopus to Crabs and Other Figures Before and After Training. The Journal of Experimental Biology, 33(4), 709–729. 10.1242/jeb.33.4.709

