## Supplementary figures and images for "Behavioral Repertoire of lab-reared early juveniles of the Mexican four-eyed octopus: *Octopus maya*"

### Supplementary figure1 and 2

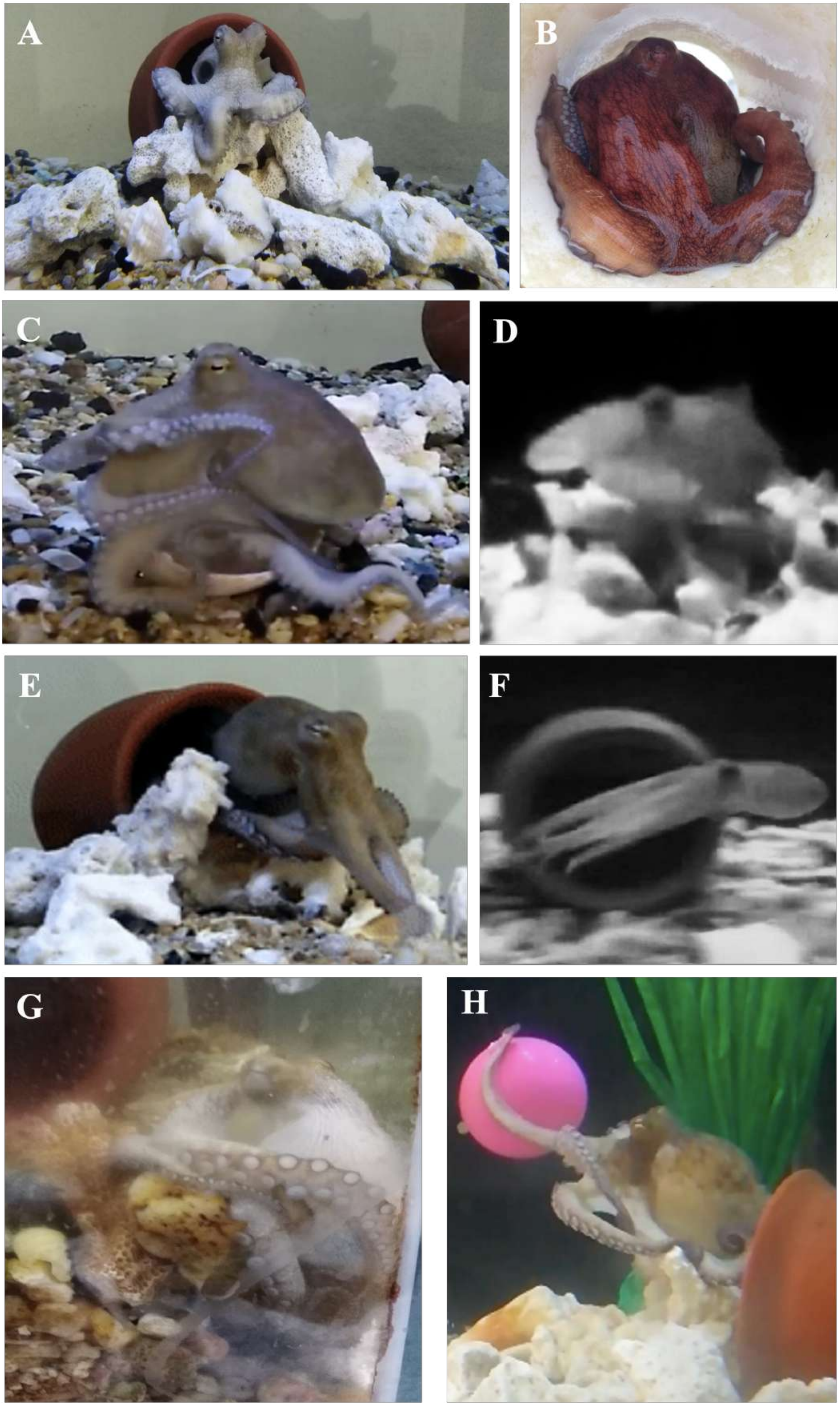

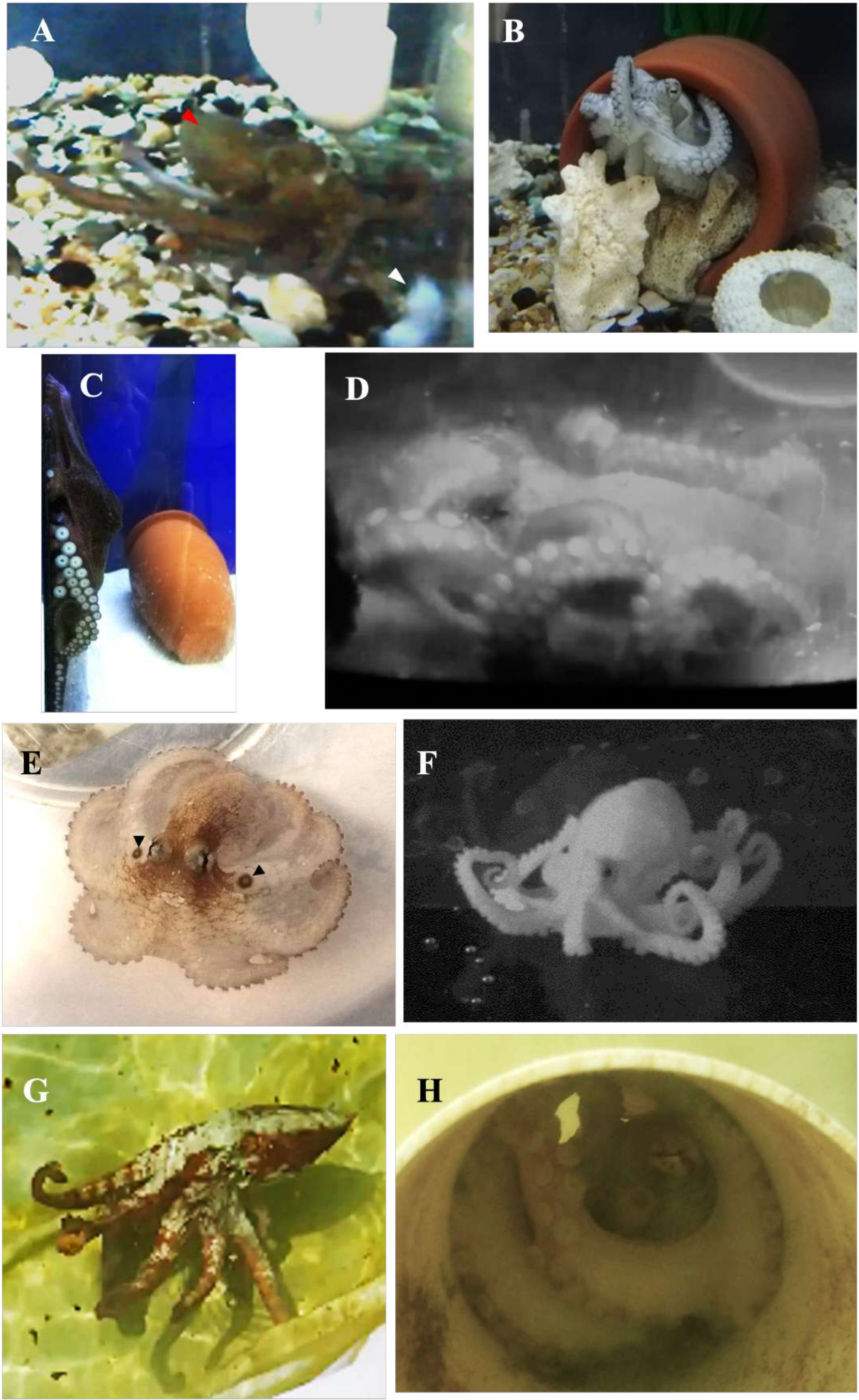
